# Quantifying the seed sensitivity of cancer subclonal reconstruction algorithms

**DOI:** 10.1101/2024.02.05.579021

**Authors:** Philippa L. Steinberg, Lydia Y. Liu, Anna Neiman-Golden, Yash Patel, Paul C. Boutros

**Affiliations:** Department of Human Genetics, University of California, Los Angeles, Los Angeles, CA, 90095, USA; Jonsson Comprehensive Cancer Centre, University of California, Los Angeles, Los Angeles, CA, 90024, USA; Institute for Precision Health, University of California, Los Angeles, Los Angeles, CA, 90095, USA; Department of Medical Biophysics, University of Toronto, Toronto, ON, M5G 1L7, Canada; Princess Margaret Cancer Centre, University Health Network, Toronto, ON, M5G 2C1, Canada; Department of Urology, University of California, Los Angeles, Los Angeles, CA, 90095, USA

## Abstract

**Background:** Intra-tumoural heterogeneity complicates cancer prognosis and impairs treatment success. One of the ways subclonal reconstruction (SRC) quantifies intra-tumoural heterogeneity is by estimating the number of subclones present in bulk DNA sequencing data. SRC algorithms are probabilistic and need to be initialized by a random seed. However, the seeds used in bioinformatics algorithms are rarely reported in the literature. Thus, the impact of the initializing seed on SRC solutions has not been studied. To address this gap, we generated a set of ten random seeds to systematically benchmark the seed sensitivity of three probabilistic SRC algorithms: PyClone-VI, DPClust, and PhyloWGS.

**Results:** We characterized the seed sensitivity of three algorithms across fourteen whole-genome sequences of head and neck squamous cell carcinoma and nine SRC pipelines, each composed of a single nucleotide variant caller, a copy number aberration caller and an SRC algorithm. This led to a total of 1470 subclonal reconstructions, including 1260 single-region and 210 multi-region reconstructions. The number of subclones estimated per patient vary across SRC pipelines, but all three SRC algorithms show substantial seed sensitivity: subclone estimates vary across different seeds for the same set of input using the same SRC algorithm. No seed consistently estimated the mode number of subclones across all patients for any SRC algorithm.

**Conclusions:** These findings highlight the variability in quantifying intra-tumoural heterogeneity introduced by the seed sensitivity of probabilistic SRC algorithms. We recommend that authors, reviewers and editors adopt guidelines to both report and randomize seed choices. It may also be valuable to consider seed-sensitivity in the benchmarking of newly developed SRC algorithms. These findings may be of interest in other areas of bioinformatics where seeded probabilistic algorithms are used and suggest consideration of formal seed reporting standards to enhance reproducibility.

## Background

Tumours arise through ancestral cells that have gained growth advantages through acquisition of somatic driver mutations, whose descendants accumulate further mutations [1]. Some cells are able to further outcompete their neighbors and form distinct cancer cell populations with shared mutations, termed *subclones* [1]. This process results in varying degrees of intra-tumoural heterogeneity in each tumour, which presents challenges to cancer prognosis and personalized treatment [2].

The intra-tumoural heterogeneity of tumour samples characterized with bulk DNA sequencing is typically quantified using subclonal reconstruction (SRC) approaches. SRC utilizes tumour DNA sequencing data to estimate the number and genotype of subclonal populations, as well as the phylogenetic relationships of subclone lineages through space and time [1]. SRC algorithms typically accomplish these tasks by first estimating the fraction of cancer cells that contain each somatic single nucleotide variant (sSNV) and then clustering these mutations based on their cancer cell fractions (CCFs) into subclones [3].

Many SRC algorithms have been proposed, each with different strengths and weaknesses in regard to accuracy, speed and robustness to varying technical and biological features [3, 4, 5, 6]. Most SRC algorithms are probabilistic. For example both PyClone-VI and PhyloWGS employ hierarchical Bayesian models and Markov Chain Monte Carlo (MCMC) for clustering mutations and inferring subclonal populations [6, 7]. PyClone-VI and DPClust use a Dirchlet prior, while PhyloWGS further uses a tree structured stick breaking prior and predicts the evolutionary relationships between subclones [7, 8, 9].

Since probabilistic models underlie SRC algorithms, they require an initializing seed for the generation of random numbers. Ideally, SRC algorithms would be seed insensitive and able to generate pseudorandom numbers regardless of the initializing seed. The initializing random seed should not significantly impact algorithm outcome, such as by generating more statistically-related numbers than by random chance and resulting in a biased starting cluster or tree structure. For varying reasons, few projects that employ probabilistic SRC algorithms report their seeds [10]. Further, the impact of seed selection on the accuracy and uniformity of SRC solutions has not been formally assessed. To fill this gap, we performed SRC using ten predetermined seeds across the SRC algorithms PyClone-VI, DPClust, and PhyloWGS, and evaluated the variability in their SRC solutions.

## Results

We reconstructed the subclonal architectures of fourteen head and neck squamous cell carcinoma tumours using nine pipelines, each constructed from a sSNV caller (three: Mutect2, Strelka2, SomaticSniper) [11, 12, 13], a sCNA caller (one: Battenberg) [9], and an SRC algorithm (three: PyClone-VI, DPClust, PhyloWGS) (**Figure 1**). To assess the impact of initializing seeds on the SRC algorithms, we performed SRC ten times per patient per SRC pipeline, each time initialized by one of ten predetermined seeds (51404, 366306, 423647, 838004, 50135, 628019,97782, 253505, 659767, 13142). SRC was performed in single-region mode, where reconstructions were focused on the primary tumour samples of the fourteen patients (total runs: 1260), as well as in multi-region mode, where reconstructions involved the primary and two lymph node metastasis samples of seven patients (total runs: 210). Across all pipelines, patients, and seeds, we launched a total of 1,470 reconstructions, with 96% success rate (1216/1260) for single-region reconstructions and 100% success rate for the 210 multi-region reconstructions.

**Figure 1:**
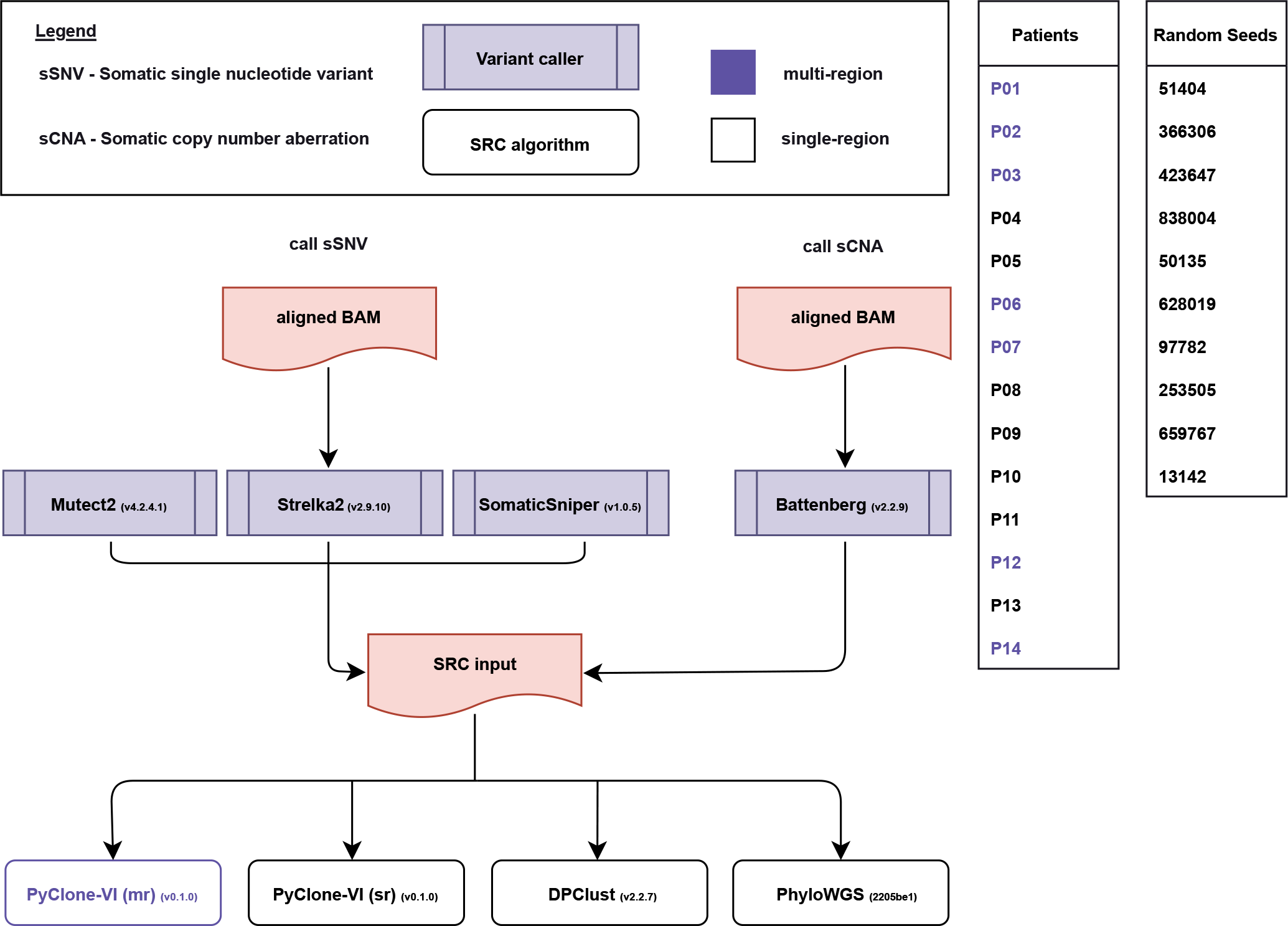
Subclonal reconstruction workflow and pipeline construction. Raw sequencing reads from paired tumour and normal samples were aligned against the GRCh38 build of the human genome using BWA-MEM and recalibrated using the Genome Analysis Toolkit (GATK). Somatic single nucleotide variants (sSNVs) were detected using Mutect2, Strelka2, and SomaticSniper. Somatic copy number aberrations (sCNA) were detected using Battenberg. Single-region primary tumour subclonal reconstruction was performed using nine pipelines combining one of the sSNV callers, the sCNA caller Battenberg, and one of the subclonal reconstruction (SRC) algorithms: PyClone-VI, DPClust, and PhyloWGS. Multi-region tumour subclonal reconstruction on one primary and two lymph node metastasis lesions (purple) was performed using the three PyClone-VI pipelines.

We assessed the seed sensitivity of the SRC algorithms and pipelines by focusing on the number of subclones detected, a metric reflective of the degree of intra-tumoural heterogeneity (**Supplementary Table 1**). Past studies have shown that sSNV caller choice impacts subclone number estimates more than sCNA caller choice [4]. Indeed, we observed variability in subclone count when inputs from different sSNV callers were used with the same SRC algorithm. Moreover, we observed variability in the number of estimated subclones in all pipelines with fixed input and SRC algorithm due to the choice of initializing seed (**Figure 2a**).

**Figure 2:**
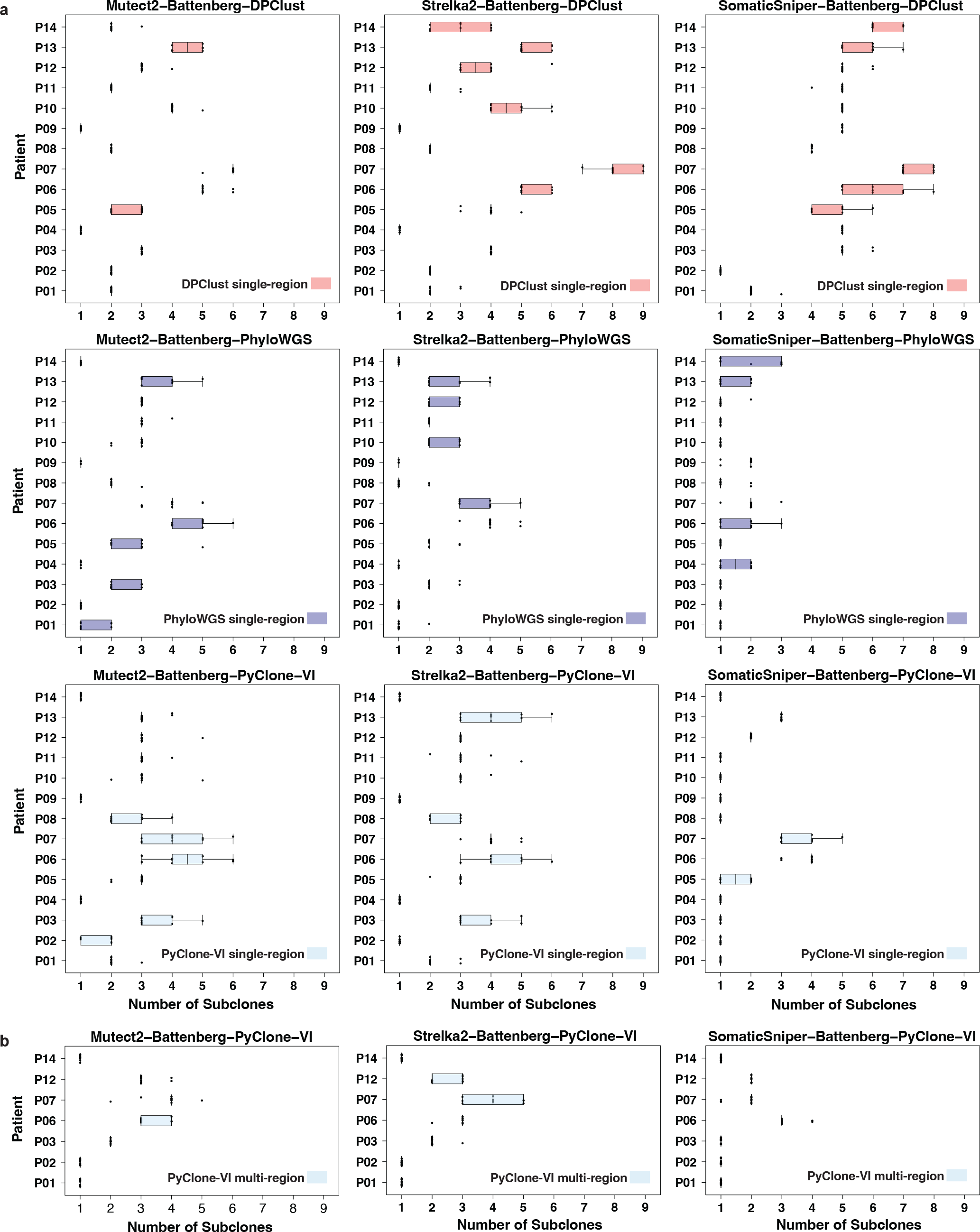
Number of subclones called across 10 random seeds. Boxplots depict median, max, 1st and 3rd quartile, and the whiskers represent maximum and minimum values within 1.5 × the interquartile range (IQR). Each dot represents a subclone number estimate initialized by a specific seed. Samples with all 10 data points on one line indicate no variation in subclone estimates across the 10 seeds. **a)** Single-region mode subclonal reconstruction (SRC) pipelines on 14 patients. **b)** Multi-region mode PyClone-VI pipelines on 7 patients.

Across single-region reconstructions, DPClust pipelines showed the greatest variability in subclone estimates across seeds (mean interquartile range [IQR]: 0.32) (**Supplementary Table 2**). This could be related to the tendency for DPClust to call more subclones (mean ± standard deviation [sd] subclone count DPClust: 3.73 ± 1.90; PyClone-VI: 2.31 ± 1.29; PhyloWGS: 1.96 ± 1.11), since in DPClust pipelines, patients with higher subclone count (median above 85th quantile) trended with higher subclone number variability due to seed (mean IQR in patients with median of <5 subclones: 0.21; ≥5 subclones: 0.50). SomaticSniper when paired with DPClust showed similar subclone number variability across seeds as when paired with PhyloWGS (mean IQR with DPClust: 0.39; PhyloWGS: 0.34), despite the uncharacteristically high number of polyclonal solutions (*i*.*e*., solutions with >1 subclone) called by the SomaticSniper-Battenberg-DPClust pipeline. On the other hand, Mutect2 when paired with DPClust showed some of the lowest subclone estimate variability across all pipelines (mean IQR: 0.14), with few to no subclone number outliers for most patients.

In contrast to DPClust, PyClone-VI showed the lowest overall single-region reconstruction variability across seeds (mean IQR: 0.28). When paired with SomaticSniper, Mutect2 and Strelka2, PyClone-VI reconstructions estimated three or more different numbers of subclones due to seed choice in only one, four and five patients, respectively (mean IQR with SomaticSniper: 0.14; Mutect2: 0.38; Strelka2: 0.32). Patients with higher subclone estimates also tended to have higher variability across seeds in PyClone-VI reconstructions (mean IQR with median <4: 0.16;≥4: 0.86). PhyloWGS had an intermediate level of consistency in estimating subclone numbers compared to the other SRC algorithms (mean IQR: 0.31), but had the most failed SRCs (44/420, 10%), compared to none from DPClust and PyClone-VI. Reconstruction failures were due to convergence on solutions with multiple primaries, consistent with previously reported results [4].

We also assessed whether seed sensitivity would differ by single-region or multi-region reconstruction when using the same SRC pipeline. Indeed, we observed higher consistency in subclone estimates across seeds for two PyClone-VI pipelines when running multi-region reconstructions (mean IQR: 0.167; with SomaticSniper: 0; Mutect2: 0.11; Strealka2: 0.43) compared to single-region reconstructions (**Figure 2b**). The differences in variability between single-region and multi-region reconstructions may also be due to multi-region solutions having lower subclone estimates on average (mean ± sd single-region: 2.31 ± 1.29; multi-region: 1.94 ± 1.10).

We further quantified the relative output variability of each predetermined seed by comparing a seed’s subclone estimate for a patient to the mode subclone estimate across the ten seeds. No seed consistently called the mode number of subclones across all patients and SRC pipelines and no seed achieved the mode number of subclones across all pipelines of any SRC algorithm (**Supplementary Table 3**). PhyloWGS had the lowest relative seed consistency of all SRC algorithms (rate of seed calling mode number of subclones: 71%), with no seed calling the mode number of subclones consistently across all patients for any PhyloWGS pipeline (**Figure 3a**). Mutect2 and Strelka2 paired with PhyloWGS also had the lowest seed consistencies out of all nine SRC pipelines (Mutect2: 64%; Strelka2: 69%). The DPClust pipelines showed a higher relative seed consistency of 81%. Mutect2-Battenberg-DPClust was still the most consistent sSNV caller and DPClust pairing, with seeds calling the mode 89% of the time and one seed calling the mode across all patients. Consistent with a low overall variability in subclone estimates, the PyClone-VI pipelines also showed the highest relative seed consistency in single-region reconstructions, with seeds achieving the mode number of subclones 82% of the time. The pipeline SomaticSniper-Battenberg-PyCloneVI showed the highest seed consistency (91%) of all single-region reconstructions, with three seeds achieving the mode number of subclones across all patients.

**Figure 3:**
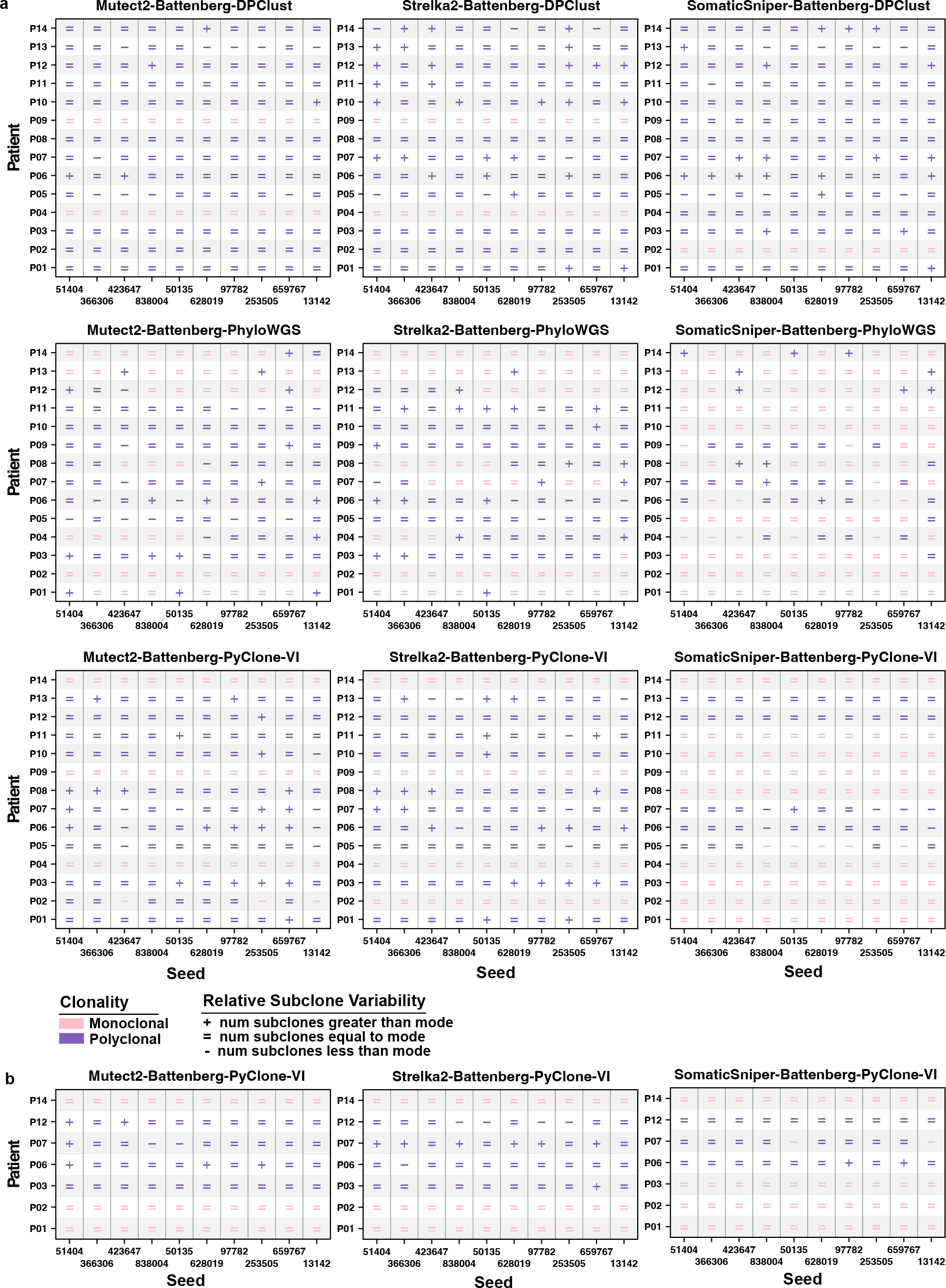
Number of subclones relative to the mode of subclones across 10 seeds. Each dot represents a subclone estimate of more than (+), equal to (=) or fewer than (-) the mode of subclone estimates per sample for an initializing seed. Pink indicates a monoclonal solution (1 subclone), and purple a polyclonal (>1 subclones) solution. Missing values indicate failed reconstruction. **a)** Single-region mode subclonal reconstruction (SRC) pipelines on 14 patients. **b)** Multi-region mode PyClone-VI pipelines on 7 patients.

The PyClone-VI pipelines in multi-region reconstruction mode also showed higher relative seed consistency (89%) than that of single-region reconstructions (**Figure 3b**). Each individual pipeline was more consistent in multi-region mode than in single-region mode, with the exception of a tie for the Mutect2-Battenberg-PyClone-VI pipeline (multi-region: 89%; single-region: 89%). SomaticSniper-Battenberg-PyClone-VI in multi-region mode had the highest rate of seeds achieving the mode subclone estimate (94%) across all single-region and multi-region reconstructions, with six seeds calling the mode across all patients. Overall, we did not observe trends of seeds consistently leading to over- or under-estimation of subclones in any SRC pipeline.

Finally, we examined the patterns of reconstruction failure in PhyloWGS pipelines in more detail. Patients P09, P04, P14 experienced the most reconstruction failures across pipelines and seeds (failure rate P09: 50%; P04: 37%; P14: 30%), while P01, P02, P11 had no failures and the rest of the patients failed rarely (3-7%) and without pattern. This suggests that differences in the underlying subclonal architecture of samples could be contributing to algorithm failure. With the overall failure rate of 10%, seeds 628019 and 50135 showed more than double the expected failure rate with 10 (24%, *p* = 0.010) and 9 (21%, *p* = 0.028) failures, respectively, though failure rates were not significantly higher than expected after multiple testing adjustment (adjusted p-value 628019: 0.10; 50135: 0.14). The seeds that succeeded across all reconstructions also showed the highest relative seed consistency in PhyloWGS pipelines (percent mode estimate per seed 366306: 85%; 42367: 83%). Taking reconstruction failure by both sample and seed influences into account, the frequent failures of some seeds (628019, 50135, 13142, 659767) but not others (366306, 42367) across all patients and sSNV callers when using PhyloWGS suggests that the seeds could be generating poor initial population structures. Thus, the choice of random seed could drastically influence SRC output, leading to outcomes ranging from variability in inferred subclone numbers to reconstruction failure.

## Discussion

Studies rarely report seeds of SRC algorithms and the impact of seed selection on SRC outcomes is not well understood. We evaluated the seed sensitivity of SRC algorithms across nine SRC pipelines each initialized by ten predetermined seeds. We found variability in subclone estimates across seeds in every SRC pipeline and no seed that consistently called the mode number of subclones for an SRC algorithm. Our results demonstrated that SRC is a seed sensitive process.

SRC algorithms are not commonly benchmarked for seed sensitivity and seed reporting rarely accompanies SRC results. This generates barriers to data reproducibility and masks any variability due to the choice of initializing seed. Studies on phylogenetic reconstruction have shown that an unspecified seed, along with unspecified parameters, processors and threads, dramatically decreases result reproducibility [14]. For PhyloWGS, some sample and seed combinations failed due to PhyloWGS finding solutions of multiple primaries, and users may inadvertently introduce bias in their reported results in the effort to find a successful seed. This underscores the importance of reporting all seeds and program configuration parameters, including those that have led to reconstruction failure.

The high variability in subclone estimates observed across seeds raises the possibility that seed selection generates systemic biases in SRC due to the inherent property of each seed. To initialize their algorithms, researchers may be choosing their starting seed manually or taking results from a pseudorandom number generator. While the goal is not to find a ‘perfect’ seed, manually chosen seeds may be too low-complexity and more stochastically dependent [15]. Further, pseudorandom number generators from standard libraries of software such as Excel, MATLAB, Mathematica, Java, python, R often do not meet quality checks for randomness [16]. To minimize seed bias, we instead recommend using the terminal to read four bytes from the operating system (/dev/random) to generate unpredictable 32-bit random seed values [17].

Further work is needed to develop statistical methods for benchmarking the influence of seeding on the probabilistic models underlying SRC algorithms. Ground truth simulations could also be applied to study how the influence of seed selection interacts with sample subclonal composition and read depth [1]. While we found convincing evidence of variability in subclone estimates due to seed in a modest cohort of 14 patients, our study should be replicated in larger cohorts to more quantitatively evaluate how each SRC algorithm is empirically affected by seed choice, and how that’s influenced by the number of regions used in reconstruction and the choice of mutation detection tools. Further, subclone count is only one of the informative features of intra-tumoural heterogeneity. To develop a fuller picture, our approach could be extended to quantify variability due to seed in other aspects of SRC, including but not limited to the cancer cell fraction estimates and mutation composition inferences of each subclone. Consequences of algorithm seed sensitivity on the potential use of intra-tumoural heterogeneity as a biomarker also need to be elucidated [18].

Beyond SRC, bioinformatics applications such as DNA aligners (*e*.*g*., BWA-MEM) [19], variant callers, and broadly, machine learning models can depend on random number generation, making them potentially seed sensitive. Our findings of seed sensitivity in SRC algorithms and pipelines may serve as the beginning of a larger discussion around seed reporting and seed sensitivity.

## Conclusions

We evaluated the seed sensitivity of three SRC algorithms by quantifying the variability in subclone estimates across ten seeds for fourteen patients with head and neck squamous cell carcinoma. We show that SRC is a seed sensitive process, as the number of estimated subclones differ across the ten initializing seeds evaluated in every pipeline. This widespread seed sensitivity makes it difficult to determine the best estimate of subclone number per patient, and could complicate the reproducibility and accuracy of SRC. Our results urge for the reporting of seed selection for all SRC tasks, and draws awareness to the under-studied problem of seed sensitivity in bioinformatics applications.

## Methods

### Aim and setting of study

The seed sensitivity of three SRC algorithms was assessed by performing repeated SRC with ten different predetermined seeds. Each SRC pipeline was composed of three algorithms: 1) a somatic single nucleotide variant (sSNV) caller, 2) a somatic copy number aberration (sCNA) caller, and 3) an SRC algorithm. The three sSNV callers, Mutect2 (v4.2.4.1), Strelka2 (v2.9.10), and SomaticSniper (v1.0.5), the sCNA caller Battenberg (v2.2.9), and the three SRC algorithms PyClone-VI (v0.1.0), DPClust (v2.2.7), and PhyloWGS (v2205be1) were combined into nine different SRC pipelines. Each SRC pipeline was run ten times per patient, each time with a different initial seed.

### Characteristics of participants or description of materials

SRC was conducted on tumour samples from an internal cohort of fourteen patients with head and neck squamous cell carcinoma (HNSC). All fourteen patients in the cohort have a primary tumour sample, but only seven of the fourteen patients have a primary tumour and two lymph node metastasis samples of sufficient quality (patients with metastasis samples = P01, P02, P03, P06, P07, P12, and P14). For single-region mode, the SRC pipeline was run on the primary tumour of each patient (n = 14). For multi-region mode, the SRC pipeline was run on the one primary and two nodal samples of each patient (n = 7).

### Clear description of all processes, inventions or description of materials: Choosing random seeds

An initial seed (3058353505) was generated from random bytes with the command *head -c 4 /dev/urandom* | *od -An -tu4* on a Linux terminal. This initial seed was used to generate ten pseudo-random seeds using *random*.*seed(*3058353505) and *random*.*sample(range(0, 1000000), k=10)* from the Python *random* library (v3.8.2). These seeds (51404, 366306, 423647, 838004, 50135, 628019, 97782, 253505, 659767, 13142) were systematically used for the SRC pipeline runs.

### Somatic variant calling

sSNV calling on paired tumour/normal BAMs was executed using the following somatic SNV callers: Mutect2 (v4.2.4.1), Strelka2 (v2.9.10), and SomaticSniper (v1.0.5) according to previously described methods [4]. Battenberg (v2.2.9) based on ASCAT (v.2.5.2) was used to perform subclonal copy number calling, according to previously described methods [4].

### Subclonal reconstruction

Three SRC algorithms were executed using one of the ten random seeds per pipeline run. PyClone-VI (v0.1.0) was executed using default settings. DPClust (v2.2.7) was executed with default settings except with no default filtering (--min_frac_muts_cluster = -1). PhyloWGS (v2205be1) was executed with parameters set to one chain (for one seed), 1000 burnins, and 2500 MCMC iterations.

### Extracting subclone output

For PyClone-VI (v0.1.0) the number of unique cluster IDs was counted as the number of subclones. For DPClust (v2.2.7) the number of unique cluster IDs in the bestClusterInfo output file was counted as the number of subclones. For PhyloWGS (v2205be1) each tree output json file was pre-processed by extracting the best tree with the largest log likelihood, reordering nodes and counting the number of nodes [18].

### Data Visualization

Figures were generated using R (v3.5.3), BoutrosLab.plotting.general (v5.9.8) [20], and Adobe Illustrator (v27.4.1). Boxplots show the median (center line), 25th and 75th percentiles (box limits), and whiskers ranging between the minimum and maximum values within 1.5 times the IQR (Tukey boxplots).

### Types of statistical analysis, power analysis

#### Data Analysis

Interquartile range (IQR) of subclone counts was calculated per patient per pipeline and averaged for pipeline and SRC algorithm arithmetic means. A high subclone median was determined as higher than the 85th percentile of subclone medians of a pipeline (PyClone-VI single-region: 3.85; PyClone-VI multi-region: 3; DPClust: 5; PhyloWGS: 3). No power analysis was performed for this study.

### Availability of data and materials

Raw sequencing files and variant calling results of the cohort used in this study are available at dbGaP, accession phs003211.v1. Analysis code can be found on the project GitHub repository https://github.com/uclahs-cds/project-method-AlgorithmEvaluation-BNCH-000082-SRCRNDSeed.

## List of abbreviations

*SRC:*: Subclonal reconstruction
*sSNV:*: Somatic single nucleotide variant
*sCNA:*: Somatic copy number aberration
*CCF:*: Cancer cell fraction
*MCMC:*: Markov Chain Monte Carlo

## Acknowledgements

The authors thank the members of the Boutros lab for their support, and the Bruins-In-Genomics (B.I.G.) Summer Program (2022) for granting ANG and PS the opportunity to work at the Boutros lab.

## Ethics declarations

## Approval and consent to participate

Not applicable.

## Consent for publication

Not applicable.

## Authors’ contributions

## Contributions

These authors contributed equally: PLS and LYL. LYL and PCB initiated the study. PLS and ANG performed subclonal reconstruction. YP developed the pipelines and helped perform experiments. PLS, ANG, and LYL conducted analysis. PLS and LYL wrote the first draft of the manuscript. LYL and PCB supervised research. All authors read and approved the manuscript.

## Supplementary Table Legends

**Supplementary Table 1: Subclone estimates per sample, seed, pipeline**. Number of subclones called across 1470 reconstructions, with 44 failed PhyloWGS reconstructions indicated with NA. sr: single-region; mr: multi-region.

**Supplementary Table 2: Subclone estimate statistics per patient**. Median, interquartile range (IQR), mean, standard deviation (sd) of number of subclones called per patient per pipeline across seeds. sr: single-region; mr: multi-region.

**Supplementary Table 3: Percent mode estimates per seed**. Percent of times that a seed called the mode number of subclones per patient per pipeline, where the mode was calculated across all seeds. sr: single-region; mr: multi-region.

